# Robust expression of transgenes in *Drosophila melanogaster*

**DOI:** 10.1101/2022.10.30.514414

**Authors:** Peter V. Lidsky, Sergey E. Dmitriev, Raul Andino

## Abstract

*Drosophila* is a classic experimental system used in fundamental and biopharmaceutical research. Manipulating gene expression in the larvae and adult flies can facilitate basic and translational studies. Here we report a method for robust transgene overexpression in *Drosophila melanogaster.* This approach is compatible with the UAS/Gal4 gene expression system. The improved expression involves a gene expression cassette that prevents transgenic mRNA transcript degradation by the nonsense-mediated RNA decay pathway. The protection is mediated by CrPV and DCV IRES sequences that apparently encode cryptic polyadenylation sites and NMD inhibitors. Stabilization of the transgene mRNA results in a >40-fold increase in its levels.

## Introduction

Controlling expression levels in the *Drosophila* model is critical for scientific and biotechnological applications^1–3^. For instance, expressing brighter fluorescent proteins is critical for imaging reporting in real-time phenotypes. Expression levels of the transgenic proteins may provide a larger dynamic range, which could better document complex phenotypes. Insect cells are widely used for protein production^4–6^. Increasing interest in insect models for pharmacological and toxicology applications. In addition, the food industry anticipates genetically modified animals can be used to produce nutritional additives or bioactive proteins^7–9^.

Not only the levels of expression are of great interest. Tools allowing production two or more proteins at once are often required. Currently, the primary approach for protein co-expression in *Drosophila* is virus-derived T2A self-processing cassettes^10^. This type of expression cassettes contains *cis*-acting hydrolase element that enable resolution of the fusion proteins by a mechanisms called ribosome skipping^11^. The T2A strategy generates proteins that have additional aminoacids at the N- and C-termini of the expressed protein^10^. Alternatively, viral internal ribosomal entry sites (IRESes) in mammalian systems are equally popular^12–17^. While IRESes are described in insect viruses^18,19^, they have never been developed into protein tools for protein production. Moreover, IRESes often lead to variable levels of the expressed proteins ^20,21^. In the context of this study we found that cryptic polyadenylation signals localized in the intragenic IRESes of dicistroviruses. This leads to truncated mRNA that do not expressed the second protein in the bicistronic construct. We found that these cryptic polyadenylation sites, when placed into standard pUAST-based^22^ constructs, result in unprecedented overexpression of the truncated transcript. Our data suggest interaction with nonsense-mediated mRNA decay (NMD) machinery as a primary reason for the overexpression and indicates the presence of in cis NMD inhibitors within the intergenic IRESes of cricket paralysis virus (CrPV)^23^ and drosophila C virus (DCV)^24^.

## Results

### CrPV or DCV IRESes result in a robust overexpression of the upstream coding region

To analyze the activity of the internal ribosomal entry sites (IRESes) *in vivo,* we engineered bicistronic constructs harboring CrPV and DCV intergenic IRESes. The first open reading frame (ORF) encoded red fluorescent protein mCherry translated via a cap-dependent mechanism, while the second IRES-directed coding region produced EGFP. Control construct encoded a mCherry-EGFP fusion protein, ensuring equimolar amounts of both reporters. The UAS promoter controlled the transcription initiation. The mRNA polyadenylation was guided by the SV40 polyadenylation signal **(Fig 1A)**. All constructs were inserted into the 3^rd^ chromosomal 86Fb-attP attachment site using ΦC31 integrase^25^. Transgenic flies were crossed with the daughterless:Gal4 (da:Gal4) driver line to test IRES-mediated expression. IRES-driven EGFP expression was not detected in most fly tissues (**Fig 1B**, middle row). In contrast, we found that the levels of mCherry produced were enormously elevated compared to the in-frame fusion control (**Fig 1B**, upper row). The level of overexpression was so high that the intense red coloration of the flies was visible in daylight with a bare eye (**Fig 1B**, bottom row).

**Fig 1.**
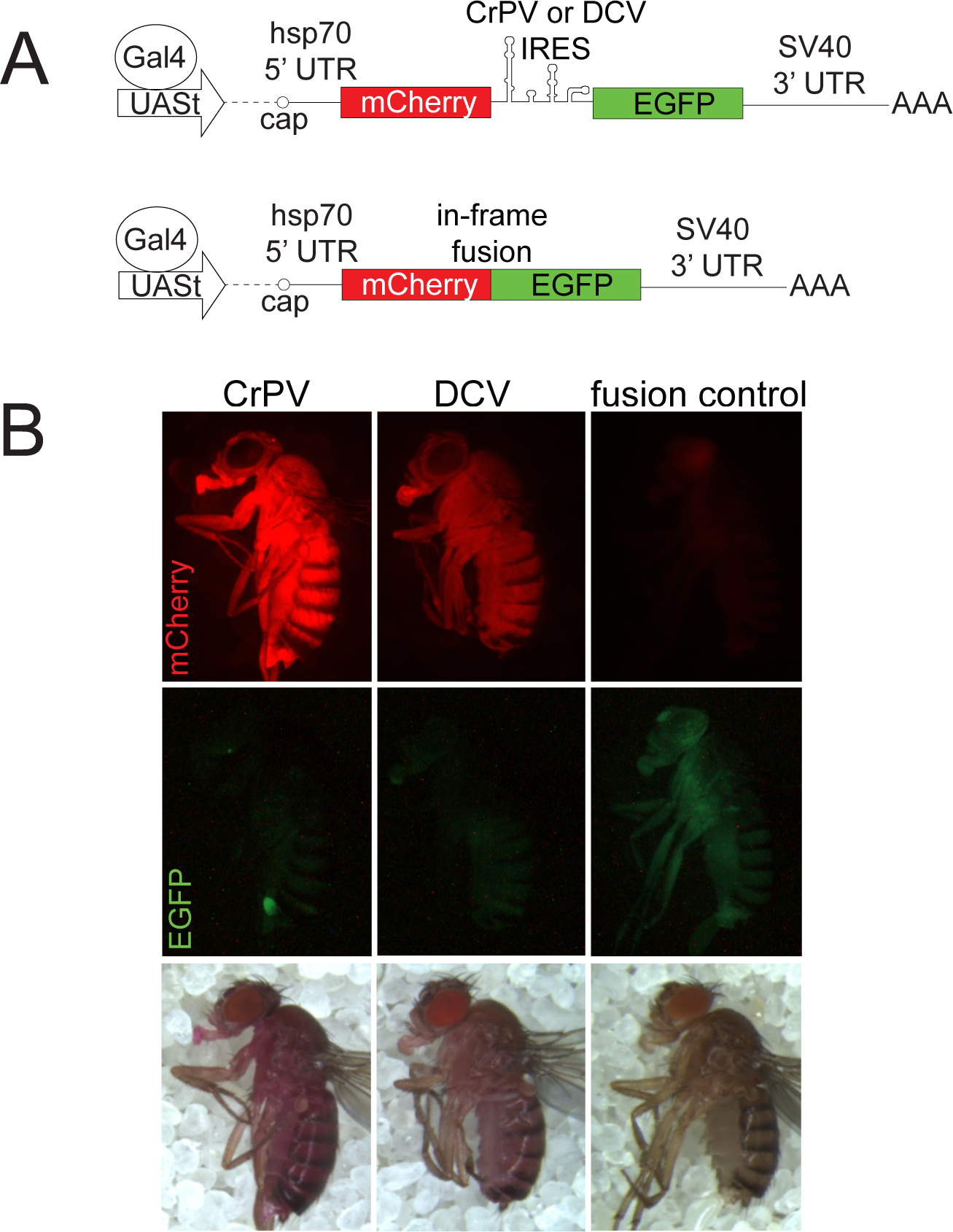
Overexpression of first cistron products in IRES-containing constructs. **A.** Scheme of the constructs used. **B.** Flies expressing *da:Gal4* and a corresponding construct (two copies of each) were examined with epifluorescent microscopy. All images are taken under the same conditions. Images in the bottom row are taken in the daylight with no excitation light.

### Levels of IRES-containing mRNAs are elevated; the transcripts are truncated and resistant to NMD

To examine the mechanism by which the first cistron is overexpressed, but the second cistron is not expressed, we examined the role of RNA decay pathways. SV40 polyadenylation cassette (~600 bp) acts as a transcriptional enhancer, and used in most UAS-based constructs. Mutations affecting nonsense-mediated RNA decay (NMD) have been reported to increase the levels of transcripts containing SV40 polyadenylation sites^26^. SV40-derived long 3’ UTRs are recognized by the NMD machinery, thus destabilizing the transcripts^26,27^. To test if NMD is responsible for IRES-mediated overexpression, we crossed *Upf2*-mutant flies with da:Gal4 and bicistronic constructs expressing mCherry and EGFP (**Fig 2A**). *Upf2*^25G^ mutation abrogates normal NMD function and is associated with late larval or pupal lethality^26^. However, flies surviving to the third instar larvae stage could be analyzed for RNA and protein expression (**Fig 2B-C**). First, we evaluated the mCherry levels with a fluorescent microscope (**Fig 2B**). The mCherry-EGFP fusion construct displayed a significant upregulation in expression in the NMD-null background (**Fig 2B,** compare fusion WT and mut). In contrast, IRES-encoding constructs did not display a significant difference in expression levels. Next, we measured the mRNA levels with the use of RT-qPCR. We used two oligo pair-sets that specifically amplify mCherry (red bars) or EGFP (green bars) coding region (**Fig 2C**). The mCherry RNA readouts were significantly higher in IRES-containing constructs compared to the fusion construct (**Fig 2C**). For instance, the RNA amounts in the IRES construct containing CrPV exceeded the fusion control RNA accumulation by more than 40-fold. Importantly, the expression of the fusion control construct was upregulated in the NMD-null background (**Fig. 2C**, compare fusion WT with mut). In contrast, CrPV construct did not display any substantial upregulation, while the DCV showed a mild, ~2 fold, increase in mCherry RNA levels. In fusion control samples, but the IRES-containing transcripts the EGFP levels are lower than those corresponding to mCherry (**Fig2C**). These results suggest that the second cistron is less stabler that the fist cistron in the bicistronic construct. Similar results were obtained with eye-specific siRNA-mediated knockdowns of NMD components *Upf1* and *Upf2* (**Fig 2D, E**).

**Fig 2.**
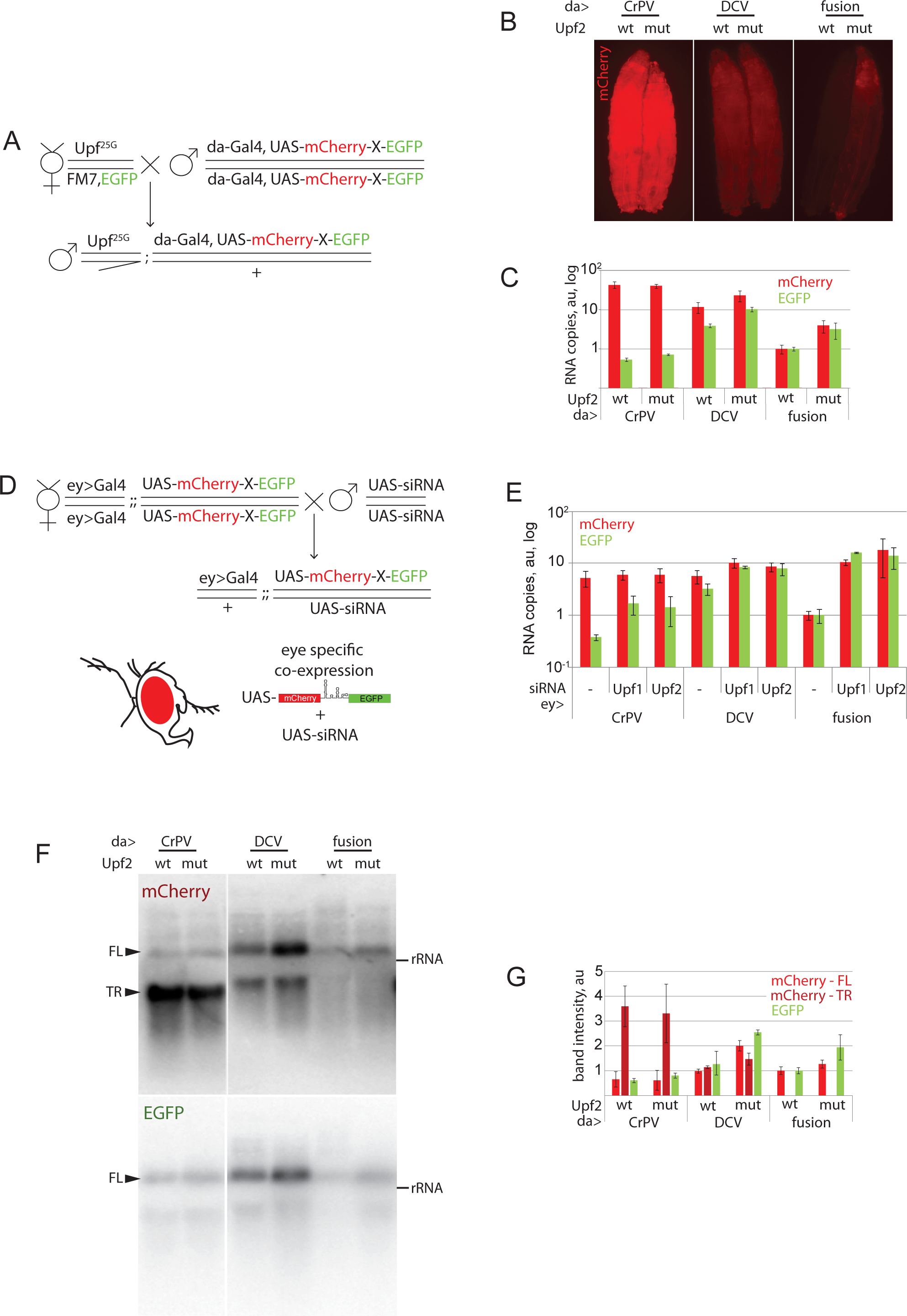
mRNAs made with IRES-containing constructs are NMD-protected. **A.-E.** Effects of *Upf^5G^* mutation only slightly affect levels of IRES-containing constructs. **A.** The crossing scheme. Hemizygous *Upf^25G^* or control males were selected for the analysis. **B.**NMD-null animals display elevated expression of the control construct but not of IRES constructs. All images are taken under the same conditions. **C.** RT-qPCR analysis of the RNA samples extracted from the animals shown in panel B. Each bar represents an average of 3 biological replicates. Numbers were first normalized with *rpl12* qPCR counts. Read-outs from wild-type animals expressing the control constructs are set as ones. A disparity in concentrations between mCherry and EGFP sequences in IRES-containing construct is evident. **D.-E.** siRNA-mediated knockdown of NMD activity only slightly affects IRES-containing constructs’ expression. **D.** A crossing scheme of eye-specific siRNA-mediated knockdown of *Upf1* and *Upf2*, the two critical components of the NMD machinery. Wild-type flies were used as control. **G.** RT-qPCR analysis performed essentially as in panel E with normalization made with Gal4 readouts instead of *rpl12.* Levels of the control construct in the wild-type background are set one. mRNAs produced with IRES-containing constructs display only a minor upregulation compared to control. An average of three independent replicates is shown. **F.** The Northern blot of the samples from panel **C**. Denatured RNA resolved in agarose gel was probed with mCherry or EGFP [α-^32^P]dCTP-labeled fragments. IRES-containing samples display truncated RNA isoform that accumulates at high levels. The full-length CrPV transcript levels were low and did not respond to NMD activity, while the full-length DCV transcript levels also accumulated to higher levels even in the presence of functional NMD. A representative image out of three replicates is shown. FL – full-length transcript, TR – truncated transcript. rRNA was used as a size marker. **G.** Quantification of band intensities from panel **F**. Each bar represents an average of 3 biological replicates. Samples from wild-type animals expressing the control constructs are set as ones.

To examine whether the IRES sequence results in RNA truncation and decay, we examine the integrity of the bicistronic construct by Northern blotting analysis using specific probes directed against either mCherry or EGFP coding sequences (**Fig 2F, G**). Consistently with the RT-qPCR results, we observed two bands corresponding to the mCherry sequences, one corresponding to the bicistronic full length transcript and the other smaller corresponding to mCherry first cistron. These shorter RNAs were apparently not sensitive to NMD. The full-length product stained with mCherry and EGFP probes was upregulated by *Upf2* inactivation in the control fusion construct but not in the CrPV IRES encoding construct. The DCV IRES containing construct showed both truncated and the full-length transcripts, which slightly increased in the *Upf2*-mutant background. Thus, we concluded bicistronic constructs (i) undergo truncation of the transcripts and (ii) provide a certain level of resistance to NMD, especially with CrPV IRES.

To determine the potential mechanism of transcript truncation, we tested if shorter transcripts (that comprise most of the mCherry-encoded RNA fraction) are polyadenylated. We purified the poly(A)+ RNAs and analyzed these with qPCR. While purification resulted in a ~200-fold decrease in the amounts of 18S ribosomal RNA (blue bars), no significant effects were found for the mCherry or EGFP-encoding RNAs (**Fig 3**). Thus, we conclude the truncated mCherry-encoding transcripts we detected are polyadenylated.

**Fig 3.**
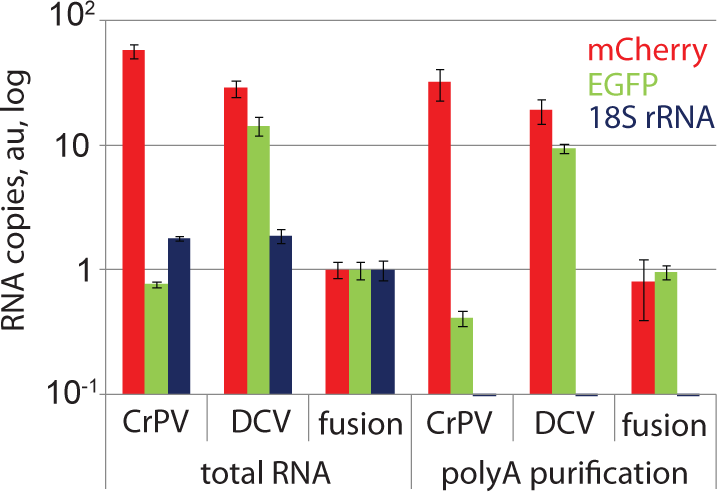
Truncated transcripts are polyadenylated since they are retained in the mixture upon isolation of the polyA fraction with the use of poly-T magnetic beads. Normalization as in panel **2C**. An average of three biological replicates is shown.

## Discussion

The experiments described above show that CrPV and DCV IRESes present in bicistronic constructs induce mRNA stabilization and a robust increase in production of the protein encoded by the 5’ proximal cistron. Based on these data, we propose the following model (**Fig 4**). In the absence of the IRESes, the SV40 polyadenylation signal plays a dual role in gene expression: first, by enhancing the transcription, second, by destabilizing the RNA by making it sensitive to NMD, in accordance with the earlier findings^26^ (**Fig 4A**).

**Fig 4.**
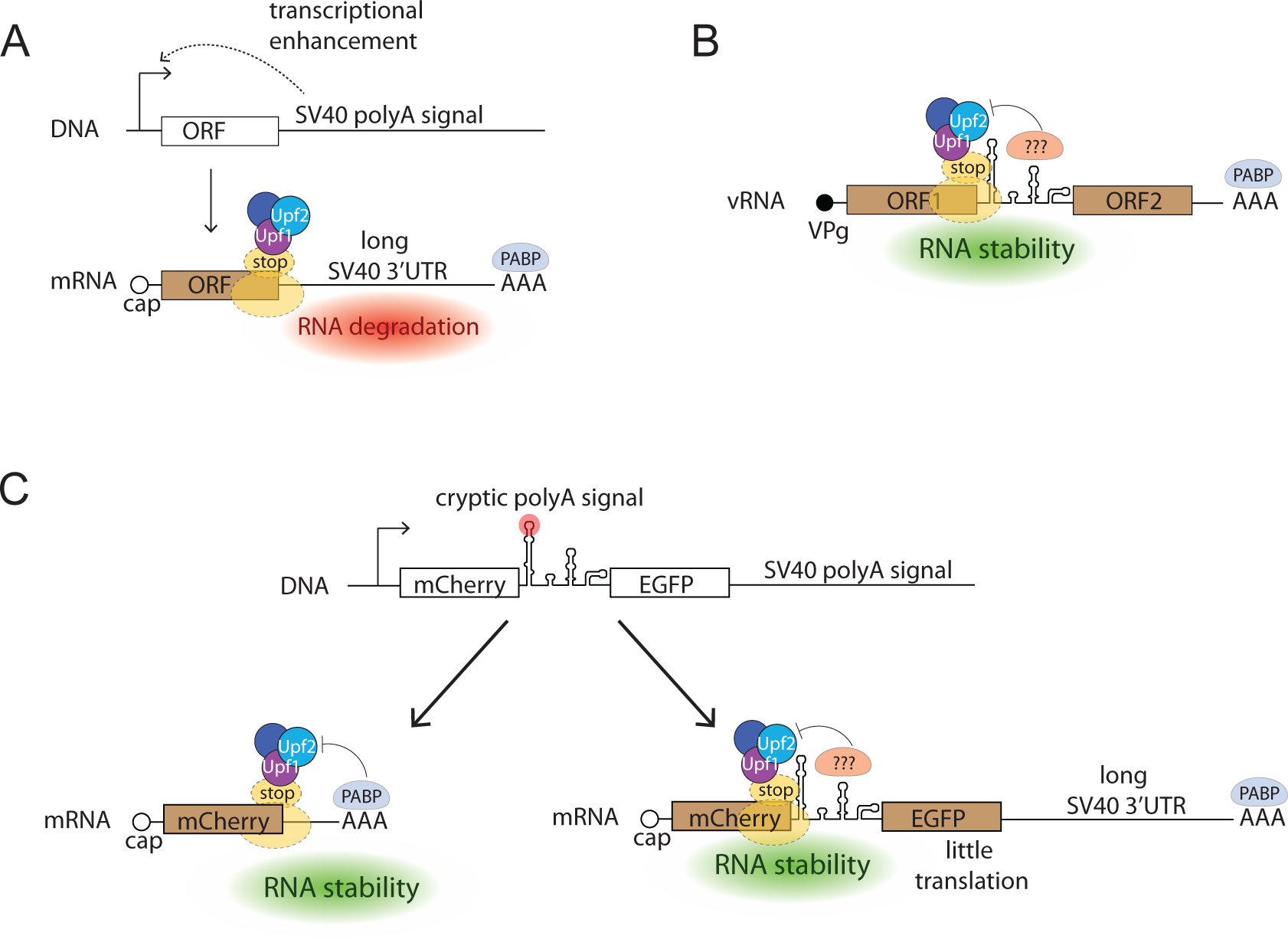
A conceptual scheme of IRES-mediated overexpression. **A.** SV40 polyadenylation signal enhances transcription but also makes the transcript sensitive to NMD. **B.** Intragenic IRES position within the virus genome provides an evolutionary rationale for the presence of NMD inhibitors within the IRESes. **C.** Cryptic polyadenylation signal within the IRES results in transcript truncation and stabilization. Putative NMD inhibitors might reside within IRESes, thus explaining decreased sensitivity to NMD inhibition.

In dicistroviral genomic RNAs, intragenic IRESes are localized immediately downstream of the first CDS encoding non-structural proteins^28^. Termination at the stop codon of this CDS should activate NMD machinery. Growing evidence suggests that NMD is also involved in the degradation of many RNA(+) virus genomes^29–33^. The NMD-inhibiting *cis*-acting RNA structures were described previously in retroviruses where these structures are located immediately after stop codons^34,35^. Unlike mammals where NMD detection relies on exon junction complex recognition, in insects NMD degrades the mRNAs that possess long 3’UTRs^36^. Termination in the first ORF of viral RNA of *Dicistroviruses* should activate the degradation pathway. The intragenic IRES sequences that take positions downstream from the stop codons might possess an additional activity to inhibit NMD (**Fig 4B**).

When placed into the intergenic spacer IRES sequences, introduce cryptic polyadenylation sites into DNA, resulting in the production of truncated transcripts that have short 3’UTR and are resistant to NMD **(Fig 2, 3)**. This retains the enhancer properties of the SV40 region but mitigates its RNA-destabilizing properties (**Fig 4C**).

In addition, the full-length bicistronic transcripts containing IRESes also displayed a decreased sensitivity to NMD if compared to the control construct. The corresponding EGFP-specific RT-qPCR readouts were not elevated (**Fig 2C**) or only slightly elevated (**Fig 2E**) in NMD-deficient strains, in contrast to those in the case of the control. Consistently, Northern blot analysis revealed that the full-length transcript in the CrPV IRES-encoding construct is not affected by *Upf2* inactivation, while the DCV IRES-encoding one is only mildly stimulated (**Fig 2C, F-G**). Thus, the full-length IRES-containing constructs also confer some resistance to NMD.

Thus, we suggest that CrPV and DCV IRESes, when put into genomic settings, might stabilize mRNA transcripts by (i) shortening the 3’ UTR due to the presense of cryptic polyadenylation signals and (ii) direct inhibition of NMD machinery.

In summary, the transcription-enhancing SV40 polyadenylation signal combined with the mRNA-stabilizing CrPV IRES constitutes a novel effective system for high-level protein production in *Drosophila*. Several approaches for tuning expression levels in flies have been published so far^37^. These methods were based on multimerization of transcription factor binding sites^37^, translational regulators^38^, or enhancers of mRNA nuclear export^39^. The novelty of our approach consists in improvement of the mRNA stability. The application of our method is not limited to fluorescent labels but could be potentially extended to other reporters and biotechnological protein production.

## Materials and methods

### Cloning and injection

To construct *pUASt-mCherry-CrPV_ires-EGFP-attB* and *pUASt-mCherry-DCV_ires-EGFP-attB* plasmids, encoding the IRES-based stress reporters, *pUASt-attB* vector was opened with EcoRI and XbaI enzymes. Fragments encoding EGFP and mCherry were amplified by PCR. EGFP fragment was digested with BglII and XbaI. mCherry fragment was treated with EcoRI and BsrGI. IRES sequences were synthesized by Genescript and excised from their vector plasmid with BsrGI and BglII. Constructs were injected into M{3xP3-RFP.attP}ZH-86Fb docking site strain^25^ by BestGene Inc.

### Fly lines

Upf2^25G^/FM7 flies^26^ were a kind gift of Prof. Mark M. Metzstein (University of Utah). A ubiquitous Gal4 driver, *GAL4-da.G32* was on the third chromosome^40^; in the text, it is referenced as da-Gal4. *y, w, ey-Flp, Act5c>CD2>Gal4* chromosome^41^ was ascribed in the text as ey>Gal4 and was kindly provided by Prof. Hugo Stocker (ETH, Zurich). Fly stocks were kept at 18°C, while the crosses - at 25°C.

### Microscopy and image analysis

Images were obtained with Nikon SMZ1500 and Leica MZ16 F. Images were deconvolved with Helicon Focus software (HeliconSoft).

### RNA

Total RNA isolation was performed with TRI-reagent (Sigma) according to manufacturer recommendations. The poly(A)-enrichment was performed with NEXTflex™ Poly(A) kit purchased from BIOO Scientific. RT reactions were performed with Maxima RT (ThermoFisher Scientific), while qPCR reactions - with SensiFAST™ SYBR^®^ No-ROX Kit (Bioline). For oligonucleotide sequences see Table S1.

**Northern blot** was performed as described^42^. 10 ug of total RNA per sample was subjected to electrophoresis on 1% (w/v) agarose gels containing 1.1 mM formaldehyde. The RNA was transferred overnight by capillarity to an Amersham Hybond-N+ membrane (GE Healthcare Life Sciences) and covalently bound to the membrane using a Stratalinker UV crosslinker. Northern blots were hybridized with DNA probes generated by a random-primed labeling reaction with use of Amersham Ready-To-Go DNA Labelling Beads (GE Healthcare Life Sciences) supplemented by [α-^32^P]dCTP and a corresponding PCR fragment as a template. Membranes were exposed overnight to a Phosphor Imager screen at room temperature.

## Acknowledgements

S.E.D. is a member of the Interdisciplinary Scientific and Educational School of Moscow University “Molecular Technologies of the Living Systems and Synthetic Biology”.

## Notes

### Competing Interest Statement

The authors have declared no competing interest.

### Summary of Updates

Figures are placed more to the center to avoid overlap with bioarxiv captions.

